# The amyloid precursor protein modulates the position and length of the axon initial segment offering a new perspective on Alzheimer’s disease genetics

**DOI:** 10.1101/2022.01.23.477413

**Authors:** Fulin Ma, Jianquan Xu, Yang Liu, Dina Popova, Mark M. Youssef, Ronald P. Hart, Karl Herrup

**Affiliations:** Department of Neurobiology, University of Pittsburgh School of Medicine; Departments of Medicine and Bioengineering, University of Pittsburgh School of Medicine; Division of Life Science, Hong Kong University of Science and Technology; Human Genetics Institute, Rutgers University; Department of Cell Biology and Neuroscience, Rutgers University

**Author notes:** Ma, Fulin < >. Liu, Yang < >. Xu, Jianquan < >. Popova, Dina < >. Youssef, Mark < >. Hart, Ronald < >.

## Abstract

The small Aβ peptide has been hypothesized to be the main driver of Alzheimer’s disease (AD). Aβ is a proteolytic cleavage product of a larger protein, the amyloid precursor protein (APP), whose normal functions remain largely unexplored. We report here activities of the full-length APP protein that relate directly to the etiology of AD. Increasing neuronal activity leads to a rapid increase in *App* gene expression. In both cultures of mouse cortical neurons and human iPSC-derived neurons, elevated APP protein changes the structure of the axon initial segment (AIS), the site of action potential initiation. In neurons with elevated APP, the AIS shortens in length and shifts in position away from the cell body. Both changes would be expected to reduce neuronal excitability. The AIS effects are due to the cell-autonomous actions of APP; exogenous Aβ – either fibrillar or oligomeric – has no effect. The findings relate directly to AD in several ways. In culture, APP carrying the Swedish familial AD mutation (APP_Swe_) induces stronger AIS changes than wild type APP. Ankyrin G and βIV-spectrin, scaffolding proteins of the AIS, both physically associate with APP, and APP_Swe_ binds more avidly than wild type APP. Finally, neurons in the frontal cortex of humans with sporadic AD reveal histologically elevated levels of APP protein that invade the domain of the AIS, whose length is significantly shorter than that found in healthy control neurons. The findings offer an alternative explanation for the effects of at least some familial AD mutations.

**Significance:** In familial Alzheimer’s disease (AD) the linkage between the genetics of APP, the neuropathology of the amyloid plaques and the symptoms of dementia are one of the strongest pieces of evidence supporting the amyloid cascade hypothesis – a conceptualization that marks the Aβ peptide as the root cause of AD. Yet, formally, the genetics only point to APP, not its Aβ breakdown product. We report here that the full-length APP protein affects the properties of the axon initial segment and through these changes serves as a dynamic regulator of neuronal activity. We propose that this newly discovered APP function offers a different, Aβ-independent, view of the genetic evidence.

## Introduction

Alzheimer’s disease (AD) is the most common cause of late-onset dementia. Most models of AD pathogenesis focus on a small peptide known as β-amyloid (Aβ) — the main constituent of the plaques found in the AD brain. The Aβ parent protein, the amyloid precursor protein (APP), was identified over 30 years ago (Goldgaber et al., 1987; Kang et al., 1987; Tanzi et al., 1987). Subsequently, mutations near the cleavage sites that release Aβ from APP were found to act as autosomal dominant AD disease genes. This linkage between the genetics of APP, the neuropathology of the plaques and the symptoms of dementia is one of the strongest pieces of evidence supporting the hypothesis that Aβ is a root cause of AD.

Formally however, the genetics link APP to AD, not Aβ. And curiously, despite the extensive research into the production and actions of Aβ, far fewer studies explore the normal functions of APP. APP is a Type I membrane protein found on cytoplasmic and vesicular membranes. It is an evolutionarily ancient protein with analogs in creatures from nematodes and fruit flies to humans (Zheng and Koo, 2006). APP is also ubiquitous, expressed in tissues throughout the body (Uhlen et al., 2015). In brain, it is found in astrocytes and neurons, but is just as abundant in oligodendrocytes and endothelial cells (Zhang et al., 2014).

APP expression begins at the pre-implantation stage of development (Fisher et al., 1991), and several developmental roles have been suggested. For example, during CNS development, mis-regulated expression of APP leads to defective cortical neuron migration (Young-Pearse et al., 2007). APP is found at the growth cone and mediates axon guidance (Kibbey et al., 1993; Rama et al., 2012; Wang et al., 2017). As neurons begin maturation, APP helps guide synapse formation (Priller et al., 2006; Klevanski et al., 2015), and promote dendritic spine formation (Lee et al., 2010). In the mature organism, APP mediates many fundamental cellular processes (Muller et al., 2017). The large extracellular domain of APP, known as secreted APP (sAPP), has a variety of functions (Mattson et al., 1993; Furukawa et al., 1996; Mockett et al., 2017). The intracellular portion of APP (known as the **A**PP **I**ntra-**C**ellular **D**omain or AICD), similar to the homologous portion of the Notch protein, translocates to the nucleus, although its function there is not well understood (Cao and Südhof, 2004). Finally, APP is a damage response protein. Multiple studies have shown that in response to stress, the levels of APP in neurons increase, particularly at a site of axonal injury (Gentleman et al., 1993; Sherriff et al., 1994; Miyai et al., 2021; Muñoz et al., 2021).

We report here on a new biological function for APP that casts many of these observations in a new light. We find that APP is a dynamic modulator of the axon initial segment (AIS), a specialized cellular compartment found at the junction between the neuronal soma and axon. It is the site of action potential initiation as well as a barrier that helps maintain neuronal polarity (Colbert and Johnston, 1996; Jenkins and Bennett, 2001; Hedstrom et al., 2008; Kole et al., 2008; Ogawa and Rasband, 2008; Nelson and Jenkins, 2017; Huang and Rasband, 2018). Through changes in its length and cellular location the AIS modulates firing probability and thus can fine tune neuronal activity (Song et al., 2009; Grubb and Burrone, 2010b; Kuba et al., 2014). We show here that upon an excitotoxic challenge, APP levels rise in the AIS, concurrent with changes in its length and cellular location. We further show that APP overexpression alone is sufficient to alter AIS position and length. Finally, we show that in APP overexpressing transgenic mice, as well as in human sporadic AD, the AIS changes are independent of the distribution of Aβ plaques. These findings offer a possible new connection between the symptoms of AD and the normal function of APP.

## Materials and Methods

### Human samples

Postmortem brain tissue was obtained from the National Institutes of Health NeuroBioBank at The University of Maryland, Baltimore, MD, the Human Brain and Spinal Fluid Resource Center, CA, and the Mount Sinai/JJ Peters VA Medical Center NIH Brain and Tissue Repository (NBTR), with approvals from Tissue Access Committee of NeuroBioBank. Subjects with Alzheimer’s disease (AD) and age-matched normal controls (NC) were requested; any cases with more than 10-hour postmortem interval were excluded. A total of 24 subjects meeting the search criteria was obtained. All tissues were from left frontal cortex (Brodmann Area 9) that had been frozen without prior fixation and stored at −80°C. Tissues were cryosectioned at 10 µm and stored at −80°C until use.

### Animals

A colony of R1.40 transgenic mice (B6.129-Tg(APPSw)40Btla/Mmjax) was established from mice originally purchased from The Jackson Laboratory (Bar Harbor, ME). The R1.40 colony is maintained on the C57BL/6J background. Wildtype littermates served as controls. Initial experiments were conducted in Hong Kong. These were conducted with under regulations established by the Government of Hong Kong SAR, and were approved by the Animal Ethics Committee of The Hong Kong University of Science and Technology (HKUST). Colonies were maintained and bred in the Animal and Plant Care Facility (APCF) at HKUST and all animal procedures complied with the guidelines of the institution and the Department of Health, Government of Hong Kong.

Additional animals used in this study were derived from stock maintained and bred in Pittsburgh in the facility maintained by the Division of Laboratory Animal Resources (DLAR), University of Pittsburgh, School of Medicine. Protocols were approved by the Institutional Animal Care and Use Committees of the University of Pittsburgh. Animals were treated in compliance with the Institute for Laboratory Animal Research of the National Academy of Science’s Guide for the Care and Use of Laboratory Animals.

Genotyping for the R1.40 transgene locus was done with the recommended PCR primers using the PCR Ready Mix kit (E3004; Sigma-Aldrich). The sequences of the genotyping primers were as follows: R1.40 Transgene Forward: 5’-CTT CAC TCG TTC TCA TTC TCT TCC A-3’; R1.40 Transgene Reverse: 5’-GCG TTT TTA TCC GCA TTT CGT TTT T-3’; Internal control forward: 5’-CAA ATG TTG CTT GTC TGG TG-3’; Internal control reverse: 5’-GTC AGT CGA GTG CAC AGT TT-3’.

### Primary culture

The culture surface of each well was coated with poly-L-lysine (0.5 mg/ml, diluted in boric acid buffer, Sigma). Wild type E16.5 C57BL/6J embryos were extracted, the cerebral cortices were isolated, and stored in chilled phosphate buffered saline (PBS) with 1 mg/ml glucose. Cortices were cut into small pieces using forceps and treated with 0.25% trypsin-EDTA (Sigma) for 10 minutes at 37 °C. The homogenized tissue was then washed in10% FBS in DMEM to inactivate the trypsin. The tissue was washed for a last time in Neurobasal medium (Life Technologies) and triturated to produce a single cell suspension. Cells were plated onto poly-L-lysine coated coverslips at 48,000 cells/well. The medium to maintain the neurons was Neurobasal medium supplemented with 2% B27 (Life Technologies), 1% Glutamax (Life Technologies), and 10,000 U/ml penicillin/ streptomycin (Life Technologies). Cultures were maintained at 37 °C in a humidified atmosphere of 5% CO_2_. Every 5 days the culture medium was refreshed by replacing half of the old medium with new.

### Human Neuron Induction

An iPS line was prepared from a healthy, 23-year-old female subject (Popova et al., manuscript in preparation). To generate glutamatergic neurons, iPS cultures were infected with lentiviruses carrying the neuronal transcription factor Neurogenin2 (*Ngn2*) and a reverse tetracycline-controlled transactivator (rtTA), in mTeSR medium (Stem Cell Technologies) with 1.5 µM ROCK inhibitor (Y27632, Tocris), as described previously (Zhang et al., 2013). The following day, the medium was replaced with Neurobasal containing B27 supplement (ThermoFisher), GlutaMAX, 2 µg/ml doxycycline and 5 µM ROCK inhibitor. Twenty-four hours following induction, transduced cells were selected with 1 µg/ml puromycin with doxycycline. Four days later, on day 5, cells were dissociated with Accutase (Stem Cell Technologies) and replated onto a monolayer of primary mouse glial cells on glass coverslips. The medium was transitioned to Neurobasal Plus medium containing B27 Plus supplement and CultureOne supplement (ThermoFisher), with GlutaMAX and 200 mM Ascorbic acid and replenished every 3–4 days. To inhibit glial overgrowth, 2 µM cytosine arabinoside was used as needed.

Treatment and morphological analyses were conducted 4–5 weeks following initial induction with doxycycline.

### Plasmids

A construct encoding human APP bearing the Swedish mutation (APP_Swe_) was kindly provided by Dr. Amy Fu of The Hong Kong University of Science and Technology. The APP-GFP construct with human APP and C-terminal GFP tag was provided by Dr. Heiman Chow at Hong Kong University of Science and Technology.

| ***Construct*** | ***Backbone*** |
| --- | --- |
| pcDNA3-APP<sub>Swe</sub> | pcDNA3 |
| pcDNA3-APP-GFP | pcDNA3 |
| pcDNA3-GFP | pcDNA3 |

### Transfection

Each plasmid was diluted to 50 ng/µL in Opti-MEM medium and incubated for 5 minutes at room temperature. Then 2 µL of lipofectamine-LTX was add to a separate 50 µL of Opti-MEM. The two portions of Opti-MEM were mixed for a final volume of 100 µL and incubated for 20 min at room temperature before adding to the neurons. Neurons were cultured in vitro for 7 days (DIV7) in a 5% CO_2_ atmosphere at a density of 24,000 cells per well of a 24-well plate with medium half-replaced once with fresh NeuroBasal medium at DIV5. The medium was changed from NeuroBasal complete to NeuroBasal without antibiotics at a final volume of 400 µL. The old, conditioned, medium was collected and stored at 4°C after which the 100 µL mixture of DNA/lipofectamine-LTX was then added without disruption into one well of the 24-well plates. The neurons were incubated for 4 hours and then changed back to their previously conditioned medium after it had been pre-warmed. The neurons were cultured at 37°C in a 5% CO_2_ atmosphere for two more days before fixation.

### Tissue processing

Animals used for collection of fresh brain tissue were anesthetized by a 1:1 mixture of 10 mg/mL Ketamine (Covetrus) and 2 mg/mL Xylazine (Covetrus) at a dose of 0.02 mL/ g. The mice were then perfused with 2 ml 0.9% saline at a flow rate of 10 mL/min. The brains were dissected free of the skull, and the cortex was dissected on the midline left hemisphere was frozen on dry ice and stored at -80 °C until use for immunoblotting. To prepare tissue for histology the right hemisphere was dissected and post-fixed by immersion in 4% paraformaldehyde for 24 hours. After post-fixation, the brains were cryoprotected by incubation in 30% sucrose in PBS for 24 hours. The brains were then embedded into OCT blocks for sectioning at 10 µm, in either the coronal or sagittal plane, on a cryostat. Sections were stored in -80°C until processed for immunohistochemistry or immunofluorescence.

### Immunochemistry

For tissue culture cells, after fixation and washing with PBS, cells were permeabilized with 0.5% Triton-X 100 in PBS (PBST). Cells were blocked for 1 hour at room temperature with PBST containing 5% donkey serum then incubated overnight at 4 °C with primary antibody in PBST at the dilutions listed in Table I. The next day, cells were washed in PBS and incubated with secondary antibody diluted at 1:500 in PBST for 1h at room temperature, followed by staining with DAPI (4’-6-diamidino-2-phenylindole) for 5 min. The coverslips were then mounted in Hydromount and viewed on a fluorescence microscope (Olympus DP80) or on a confocal microscope.

For brain tissue, cryosections were immersed in citrate buffer (pH 6.0) and heated at 95 °C for 10 mins for antigen retrieval. After cooling and rinsing, the sections were incubated in 10% donkey serum at room temperature for 1 hour to reduce non-specific antibody binding followed by overnight incubation at 4 °C with the primary antibody. The primary antibodies used, along with their sources are listed in Table I. After rinsing, the sections were incubated in Alexa Fluor conjugated secondary antibody for 1 hour at RT. Nuclei were stained with DAPI (4’-6-diamidino-2-phenylindole). Fluorescent imaging was done on a Nikon Eclipse Ti2 Confocal microscope with a 60X (1.4 Numerical Aperture) oil objective lens and A1R (Andor, 512µm × 512µm) camera, X-Cite 120 LED illumination system, and a DAPI (400-418 nm), FITC (450-490 nm), TRITC (554-570nm) and Cy5 (640-670 nm) filter sets controlled by NIS-Elements AR software.

### RNA isolation and qPCR

DIV14 neurons treated with 1 µM glutamate for the times indicated in the text then harvested using the Qiagen RNeasy Mini kit (Qiagen, 74104) according to the manufacturer’s instructions. For each sample, an equal amount (500 ng) of RNA was reverse transcribed to cDNA using iScript reverse-transcription kit (Bio-rad, 1708841). Quantitative real time PCR (45 cycles – LightCycler480 software) was performed using fast SYBR Green Master Mix (Bio-rad 1725122). GAPDH served as loading controls. The sequence of *App* primers are as follows: Forward – ACTCTGTGCCAGCCAATACC; reverse – GAACCTGGTCGAGTGGTCAG.

### Stochastic optical reconstruction microscopy

For stochastic optical reconstruction microscopy (STORM) imaging, cells were incubated with the primary antibody at 4°C overnight, and then with Alexa647-conjugated and CF568-conjugated goat-anti-rabbit/mouse secondary antibody for 2 hours at room temperature. STORM imaging was done on a custom-built STORM imaging system in the Liu laboratory based on an Olympus IX71 inverted microscope frame with a 100x, NA=1.4 oil immersion objective (UPLSAPO 100XO, Olympus), as described previously (Xu et al., 2018). For two-color STORM imaging, 30,000 frames were acquired at an exposure time of 20 ms per frame for each color (642nm and 561 nm). Online drift correction was independently performed every 200 frames (∼ 4 s) with fluorescent beads (Thermo Fisher Scientific, F8803) as fiduciary markers (excited with 488 nm laser) throughout the image acquisition process, based on our established method (Ma et al., 2017).

### Antibodies

| Target Name | Manufacturer | Catalogue Number | Dilution used |
| --- | --- | --- | --- |
| MAP2 | Abcam | Ab5392 | 1:2500 |
| 6E10 | Covance | SIG-39320 | 1:1000 |
| APP-N' | Abcam | Ab15272 | 1:400 |
| AnkG | Millipore | MABN466 | 1:500 |
| Pan-Nav | Millipore | AB-5210 | 1:500 |
| Bassoon | Abcam | 82958 | 1:500 |
| βIV Spectrin | Dr. Mathew Rasband | N/A | 1:250 |

#### Western blots

Cultured cells were harvested and homogenized in ice-cold RIPA buffer (EMD Millipore) with 1X PhosSTOP phosphatase inhibitor mixture (Roche Applied Science) and 1X complete protease inhibitor cocktail (Roche Applied Science). The homogenate was then centrifuged at 4 °C for 20 minutes at 15,000 rpm. The supernatant containing protein was collected. Protein concentration was determined using the Bradford assay (Bio-Rad). A total of 10 µg protein was then separated by SDS/PAGE and transferred to Immuno-Blot PVDF membranes (Bio-Rad). Membranes were incubated in 5% non-fat milk for 1 hour at room temperature to block nonspecific binding. Primary antibodies were applied at room temperature overnight. After rinsing three times with TBST, membranes were incubated with secondary antibodies for 1 hour at room temperature. Signals were visualized with SuperSignal West Pico, Dura, or Femto chemiluminescent substrate (ThermoFisher Scientific).

#### Co-immunoprecipitation

Wild type and R1.40 brains were homogenized in a Dounce homogenizer. The lysates were then incubated in co-IP buffer (20 mM Tris HCl (pH 8.0), 137 mM NaCl, 1% Nonidet P-40 (v/v) and 2 mM EDTA) on ice for 30 min. The brain lysates were incubated with primary antibody and protein G magnetic beads overnight at 4 °C with moderate shaking. Then the beads were washed with co-IP buffer three times and eluted with 1x SDS-PAGE loading buffer. The eluted proteins were then run on SDS/polyacrylamide gels and subjected to western blotting.

#### Aβ Fibril and oligomer preparation

Beta amyloid (Aβ_1-42_) aggregation kit was purchased from rPeptide (A-1170-2). To produce fibrillar Aβ, the Aβ powder was dissolved in NeuroBasal medium to a final concentration of 220 µM, followed by incubation for 5 days at 37 °C. To prepare Aβ oligomers, 22.2 µL DMSO was added to 0.5 mg of HFIP treated Aβ_1-42_ (A-1163-1) to a final concentration of 5 mM. The Aβ and DMSO mixture was then vortexed for 30 seconds to make sure all the Aβ powder was dissolved in DMSO followed by a short centrifugation. The solution was sonicated for 10 minutes in a bath sonicator followed by the addition of 1087 µL cold 4°C phenol-red free F-12 medium to achieve a final concentration of 100 µM. The Aβ solution was then vortexed for 15 seconds and incubated at 4°C for 24 hours. Afterwards, Aβ was centrifuged at 14,000 g for 10 minutes at room temperature. The supernatant was then collected for use.

### Statistical analysis

Data are expressed as mean ± SEM (standard error of the mean) for each group. The statistical significance of changes in different groups was evaluated by unpaired student t-test using GraphPad Prism software. P-values less than 0.05 were considered to be significant. In each figure the following code is used to describe the p-values – *: p < 0.05, **: p < 0.01, ***: p < 0.001, ****: p < 0.0001.

## Results

### Hyperactivity increases the level of APP and causes AIS length and position to change

APP is recognized as a stress response protein. Following oxidative stress, excitotoxicity, or exposure to free radicals, APP protein levels increase (Frederikse et al., 1996; Masliah et al., 1997; White et al., 1999). This is relevant to the situation in AD as excitotoxicity is cited as an important risk factor (Greenamyre and Young, 1989). Indeed, blunting the effects of excitotoxicity is the theoretical mechanism of action of the FDA approved drug, memantine (Johnson and Kotermanski, 2006). To test this association of APP with excitotoxicity in vitro, we used E16.5 mouse cortical neurons cultured for 14 days in vitro (DIV14) and exposed to a relatively low concentration of glutamate (1µM). We found that in treated, but not untreated cells, the intensity of neuronal APP immunostaining increased noticeably 24 hours following glutamate addition (Fig. 1A-D). The increased fluorescent signal was obvious in the cell body as well as in the dendrites (MAP2-positive processes) and was blocked by pre-incubating the neurons with 20 µM dCPP, a selective NMDA receptor antagonist (Monnet et al., 1995; Costa et al., 2009).

**Figure 1.**
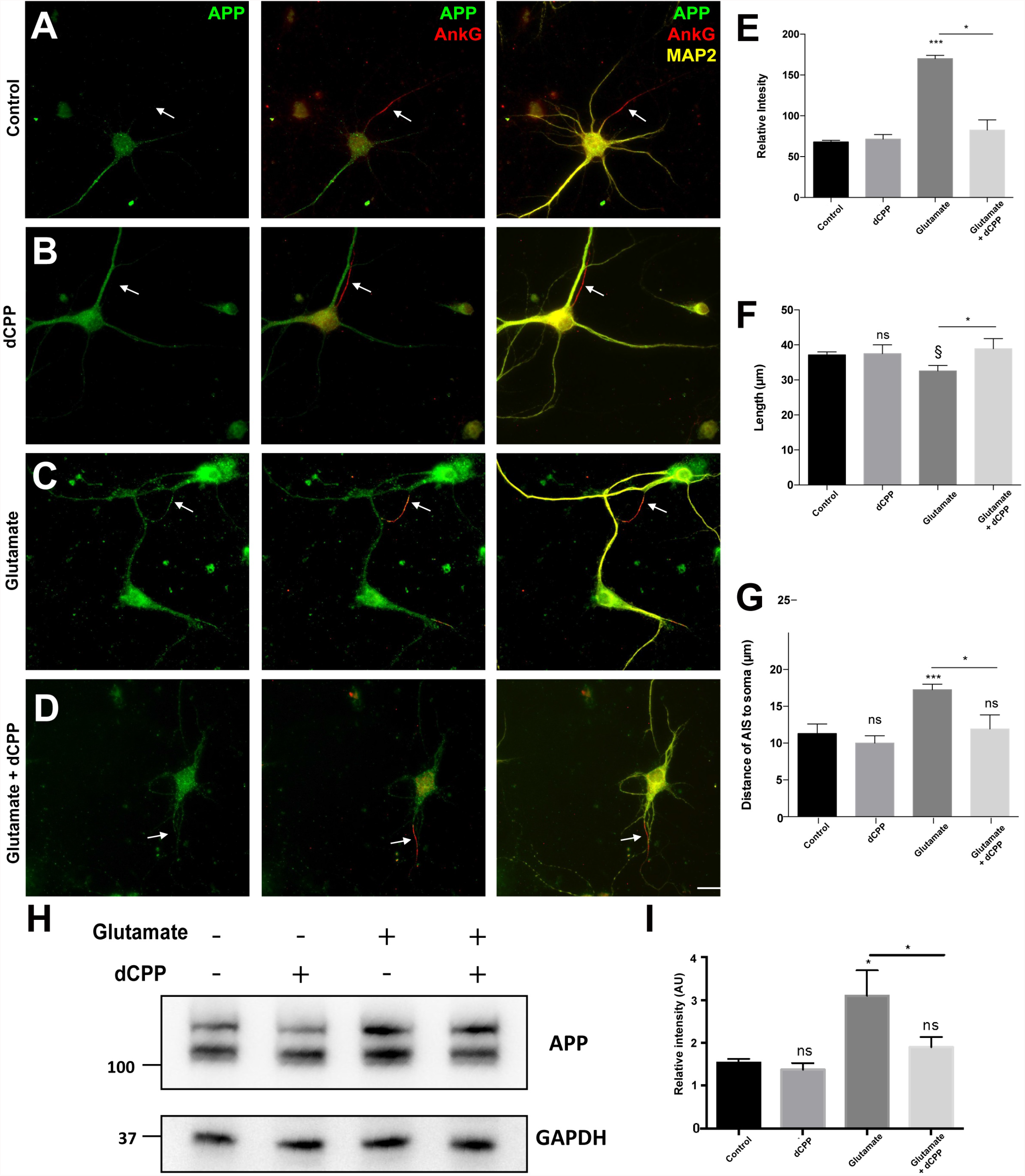
Glutamate stimulation affects cellular APP levels and the length and location of the AIS. (**A-D**) Immunofluorescent staining of DIV14 primary cultured neurons after 24-hour treatment with 1µM glutamate, with or without the protection of dCPP (APP (green), ankyrin G (AnkG, red), and MAP2 (yellow)). (**A**) untreated control; (**B)**: 20µM dCPP: (**C**): 1µM glutamate; (**D**): 20µM dCPP added 30 min before 1µM glutamate. (**E**) Intensity of APP immunofluorescent signal in AIS (defined as the AnkG-positive region). Fluorescence measured using Fiji software. (**F**) Length of the AIS after the indicated treatments. (**G**) Distance from the cell soma to the proximal end of the AIS. (**H**) Western blot of APP protein after treatments. (**I**) Quantification of band intensities shown in H. *, p<0.05; ns, §, not significant. one-way ANOVA; n=4. Scale bar, 10 µm. Data represent means ± SEM.

While the quantitative increase in staining was evident throughout the cell, we also observed a qualitative change in the axon and AIS. We identified the axonal process and the location of the AIS by immunostaining the AIS scaffold protein, ankyrin G (AnkG, red signal, Fig. 1A-D, white arrows). In untreated cells the axon was nearly devoid of APP immunoreactivity (green signal, Fig. 1A). After treating with 1 µM glutamate, however, the increased APP signal infiltrated the axon hillock and the AIS (Fig. 1C). Using ImageJ software, we found that, on average, the APP fluorescent signal intensity increased more than two-fold in the AIS of treated neurons (Fig. 1E). This expansion of the domain of APP staining was found in virtually every neuron in our cultures.

In addition to the increased level of APP in glutamate treated neurons, the properties of the AIS also changed. Its length was significantly reduced, and the distance between the cell body and the proximal end of the AIS was significantly increased (Fig. 1F, G). These changes were mediated by glutamate directly, acting through the NMDA receptor. A 30 min pretreatment of the cultures with dCPP completely blocked the AIS changes as well as the increase in APP protein (Fig. 1H, I). One plausible interpretation of these findings is that the extra activity caused by glutamate led to an increase in APP and the increased APP led to the changes in the AIS. The AIS changes in turn would be expected to increase the threshold for initiating an action potential (Grubb and Burrone, 2010b; Evans et al., 2013b; Muir and Kittler, 2014) thus making the neurons more resistant to subsequent stimuli, essentially providing protection against excitotoxicity.

### Overexpression of APP is sufficient to change the AIS configuration

The results from the glutamate experiments suggest that APP regulates the configuration of the AIS to modulate neuronal excitability. To test this hypothesis, we asked whether, in the absence of a glutamate stimulus, APP alone would be sufficient to produce similar effects. We co-transfected APP and GFP-encoding plasmids into DIV7 cultured neurons. Control cultures were transfected with a GFP plasmid only. Two days later the cells were fixed and immunostained for MAP2 and either βIV spectrin or AnkG (Fig. 2A, B). Transfected neurons had elevated APP that infiltrated all processes, including the axon. Consistently, in APP expressing neurons, the length of the AIS was reduced by almost one-half compared to control. The distance from the cell body to the proximal end of the AIS also increased (Fig. 2C). Thus, APP over-expression alone, in the absence of glutamate stimulation, is sufficient to cause the AIS to shift away from the cell body and to shorten

**Figure 2.**
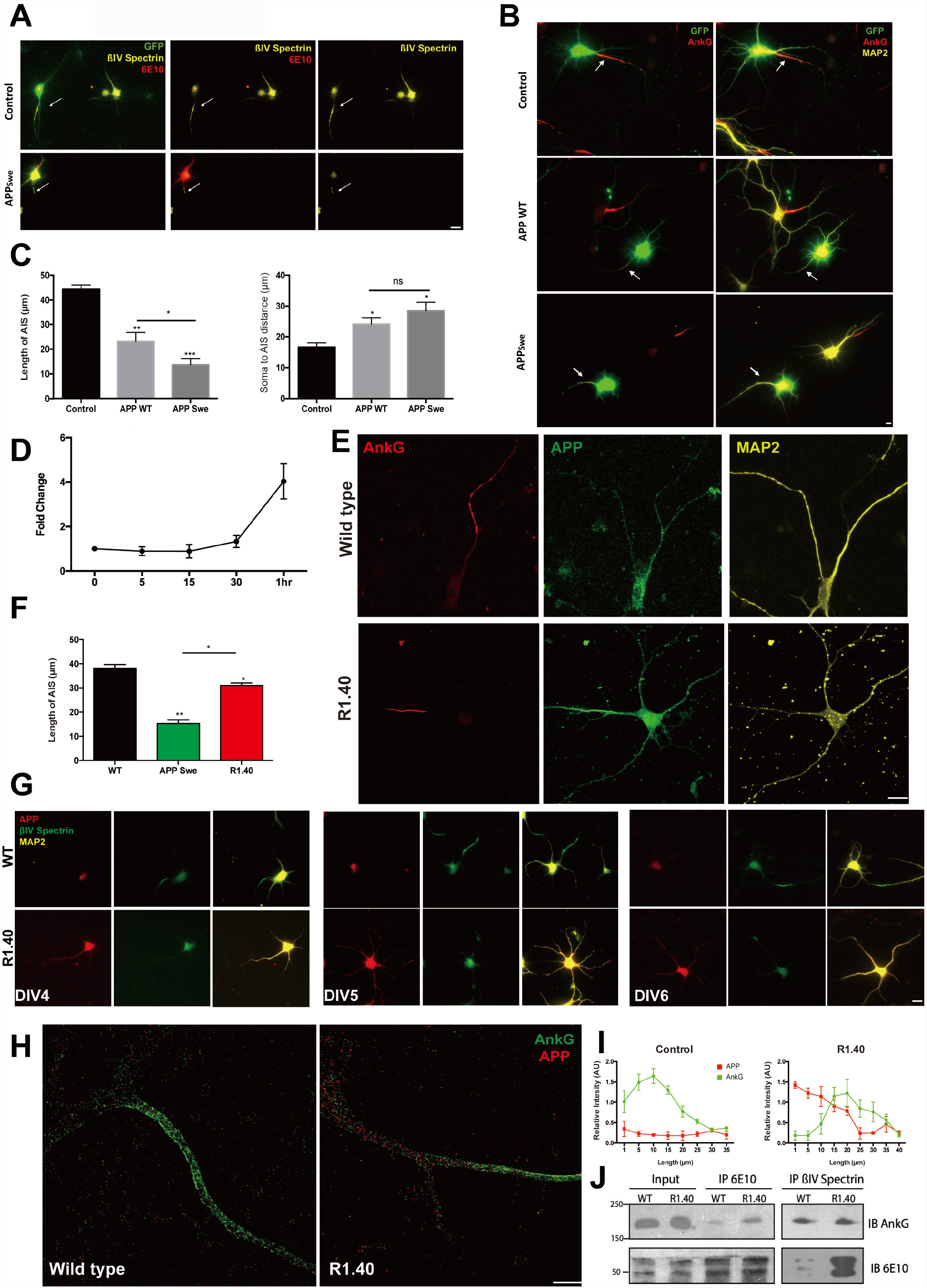
APP overexpression induces AIS changes. (**A**) Cultured cortical neurons, co-transfected on DIV7 with *APP*_*Swe*_ and *GFP* plasmids, were immunostained for βIV spectrin (yellow) and 6E10 (red) 2 days after transfection. Scale bar = 10 µm (**B**) Cultured cortical neurons were co-transfected on DIV7 with GFP as well as with either wild type *APP* (APP WT) or *APP*_*Swe*_ plasmids. Cultures were fixed and immunostained for βIV spectrin (yellow) and 6E10 (red) 2 days after transfection. Scale bar = 5 µm. (**C**) Quantification of the AIS length (left) and distance to soma (right) in transfected neurons. (*, p<0.05; **, p<0.001; one-way ANOVA; n=4. Data represent means ± SEM.) (**D**) RT-PCR measurements of *APP* mRNA at various times after treatment with 1 µM glutamate. (**E**) Confocal image of the AIS immunolabeled with AnkG (red), N-terminal APP antibody (N’APP. green), and MAP2 (yellow). Cells were from DIV14 cortical neuron cultures from wild type (top row) or R1.40 (bottom row) cultures. Scale bar = 1 µm. (**F**) Quantification of the AIS lengths after transfection with *APP*_*Swe*_ (e.g., panel B) or in cultures of R1.40 neurons. (*, p<0.05; **, p<0.001; one-way ANOVA; n=4. Data represent means ± SEM.) (**G**) Immunolabeling of APP (red) in the developing axon using βIV spectrin (green) to label the AIS. MAP2 (yellow) was used as a dendritic marker. Scale bar = 5 µm (**H**) Super-resolution imaging of the of AnkG (green) and N’APP (red) of cultured neurons from wild type and R1.40 mice fixed at DIV 14. Scale bar = 5 µm. (**I**) Intensity (relative gray value) of APP (red) and AnkG (green) in the AIS as a function of distance from the cell body. (**J**) Western blots of co-immunoprecipitation of cortical lysates from 18-mo-old wild type (WT) and R1.40 mice probed with AnkG and 6E10 antibody.

If these APP-induced changes serve to buffer the neuron against the damage caused by excessive neuronal activity, the expectation would be that the APP response would be rapid after the initial stimulus. We tested this idea by exposing DIV14 neuronal cultures to 1 µM glutamate for varying lengths of times, then harvested the cells and performed qPCR on the cell lysates. The response of *App* gene expression was rapid and robust. After 30 minutes, the level of *App* mRNA started to increase and within one hour of stimulus was increased 3- to 5-fold (Fig 2D). This level of responsiveness, combined with the effects of APP on the AIS is consistent with the hypothesis that APP serves as a buffer against neuronal hyperexcitability.

### The AIS in Alzheimer’s disease – in vitro data

That APP overexpression can result in significant changes in the AIS raises the question of whether the elevated levels of APP that are found in Alzheimer’s disease have similar effects and thus possibly contribute to the symptoms of dementia. We tested this idea first by comparing the properties of the AIS in neurons transfected either with wild type human *APP* (*hAPP*) or with the well-characterized Swedish variant – K595N/M596L (*hAPP*_*Swe*_) that is a recognized Alzheimer’s disease gene. We found that hAPP_Swe_ caused a shrinkage of the AIS that was nearly 50% greater than that found with wild type hAPP (Fig. 2B, C). Note that neither wild type hAPP nor hAPP_Swe_ caused visible damage to the transfected neurons. At two days post-transfection the appearance of the MAP2 immunostained dendritic structure was not noticeably notably different from neurons expressing only the control GFP vector.

We were concerned that the levels of APP protein after transfection would be artificially high – so much so that they would create a cellular state that did not reflect normal physiological conditions. We therefore cultured cortical neurons from brains of the R1.40 Alzheimer’s disease mouse model. The R1.40 *hAPP*_*Swe*_ transgene is expressed at only 2-3 times the levels of the endogenous mouse *App* gene (Lamb et al., 1993; Lamb et al., 1997). This is close to the increase in mouse *App* expression observed following a glutamate challenge (Fig. 2D). Even under these more physiologically relevant levels of expression, however, the response of the AIS was still significant. We found that in R1.40 neurons axonal APP immunostaining invaded the axon hillock as well as the AIS, and the length of the AIS was reduced compared to neurons cultured from wild type animals of the same litter (Fig. 2E). Compared to the *APP*_*Swe*_-transfected neurons described above, the reduction of the length of the R1.40 AIS was less dramatic (Fig 2F). This suggests that the APP-driven shortening of the AIS is both dose and genotype dependent. Intriguingly, the APP effect is seen even during neuronal maturation. We used βIV spectrin to track the appearance of the AIS as a function of time in culture. In wild type neurons, the AIS begins to form in the growing axons (monitored by βIV spectrin) after DIV4. In R1.40 neurons, by contrast, the AIS does not appear until DIV6 (Fig. 2G)

To obtain a more fine-grained picture of the interplay of APP with the components of the AIS, we used stochastic optical reconstruction microscopy (STORM). The resulting high-resolution images showed that in wild type mouse cultures, endogenous APP (stained with an APP N-terminal antibody) gave a weak signal in the AIS but was clearly visible in the cell body (Fig. 2H). In R1.40 neurons, however, APP staining was stronger and could be found entering the axon and the region of the AIS. The impression gained when comparing this pattern with that of the AnkG scaffold protein was that APP was “pushing” the AIS away from the cell body. Quantification of the relative signal intensity of the two antibodies was consistent with this idea (Fig. 2I).

We next asked whether the AIS scaffold protein physically interacted with APP. We prepared lysates from R1.40 and wild type neocortex and performed immunoprecipitations with APP, AnkG, or βIV spectrin antibodies (Fig. 2J). Immunoprecipitation of R1.40 mouse brain lysates using the 6E10 (APP) antibody pulled down AnkG (Fig. 2J, lane 4). A similar result was found with wild type lysates but the strength of the AnkG band was considerably reduced (Fig. 2J, lane 3). We used separate brain lysates to show that βIV spectrin immunoprecipitates also contained APP (Fig. 2J, lanes 5, 6). The amount of APP pulled down by the βIV spectrin antibody from R1.40 lysates was greater than from wild type despite comparable input bands (Fig. 2J, bottom gel, lanes 5, 6). One interpretation of this difference is that the interaction between the AIS scaffold proteins and the endogenous mouse APP is weaker than that between the scaffolds and hAPP_Swe_, encoded by the R1.40 transgene Taken together these results demonstrate that APP, a membrane protein implicated in Alzheimer’s disease, is intimately associated with two of the most important scaffolding proteins of the AIS.

Enhanced expression of APP protein – either wild type or APP_Swe_ – is associated with increased proteolytic processing and hence with elevated levels of the Aβ peptide in the culture medium. This raised the possibility that the effects of APP expression were indirect and mediated through Aβ. To test this idea, fibrillar Aβ [Aβ(f)] and Aβ oligomers [Aβ(o)] were prepared from commercially available peptide by standard protocols (Mehta et al., 1998; Zhu et al., 2019). The level of aggregation was verified by western blot using the 6E10 antibody as a probe (Fig. 3A). The Aβ preparations – at concentrations used previously, 5 µM Aβ(f) and 1µM Aβ(o) – were then used to treat DIV14 neuronal cultures. After 24 hours, the cells were fixed and immunostained for AIS scaffold proteins and markers of synapses. After Aβ treatment, neuronal number remained unchanged, but the density of synapses was reduced compared to untreated controls (Fig. 3B). Despite the loss of synapses, the length of the AIS remained unaffected in both the Aβ(f) and Aβ(o) treated cultures (Fig. 3C, D). The distance of the proximal end of the AIS from the cell soma likewise showed no significant change (Fig. 3E).

**Figure 3.**
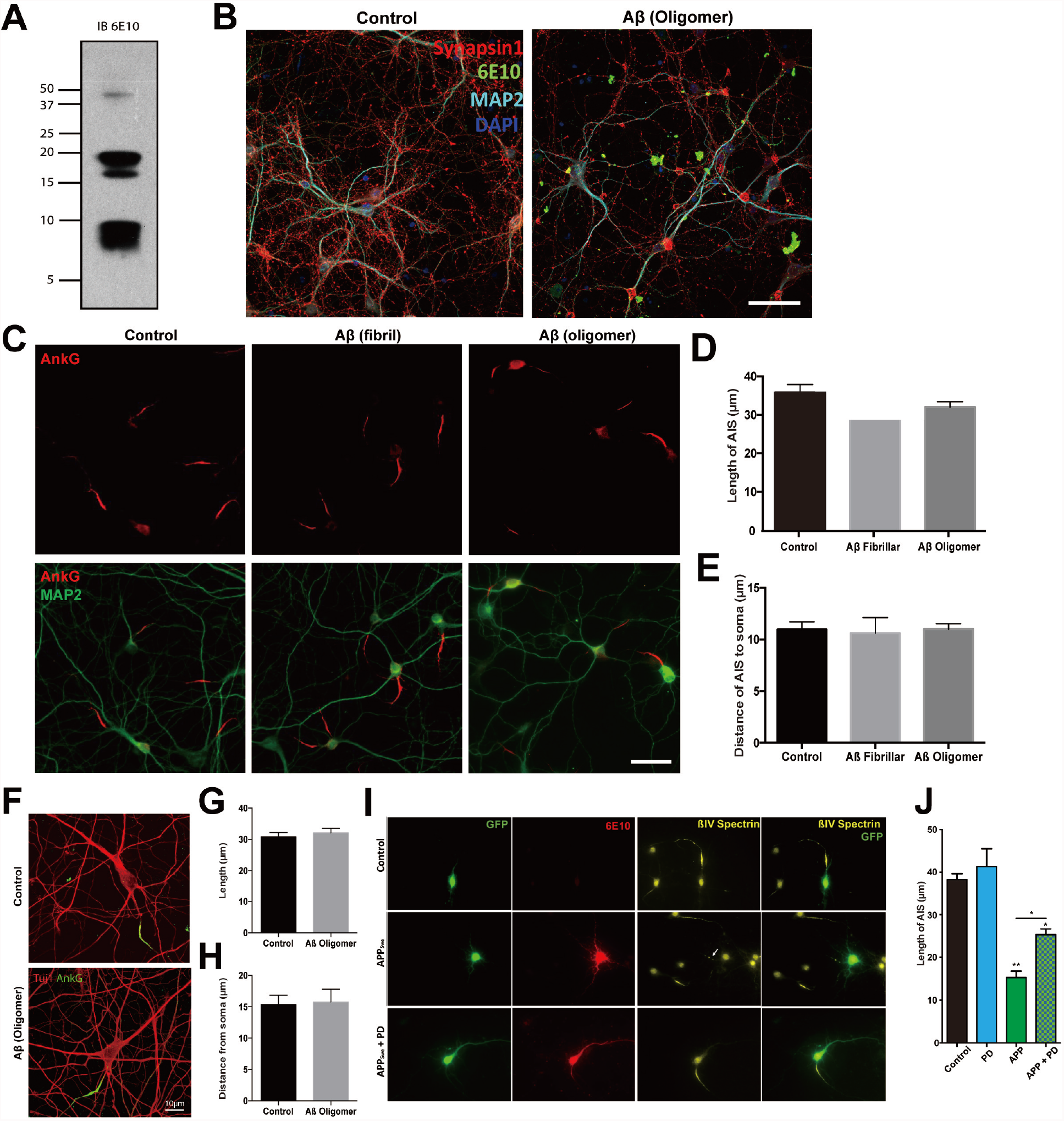
APP induced changes in the AIS changes are independent of Aβ but partially dependent on calpain activity. (**A**) Western blot of the Aβ oligomer sample used (blotted with the 6E10 antibody). (**B**) DIV14 primary cultured neurons treated with Aβ oligomer (1 µM) for 24 hours, then fixed and immunolabeled for Synapsin1 (red), Aβ (6E10 antibody, green), MAP2 (cyan) and DAPI (blue). Scale bar = 20 µm. (**C**) DIV14 neurons treated with fibrillar or oligomeric Aβ for 24 hours then immunolabeled for AnkG (red) to reveal the AIS and for MAP2 (green). Scale bar = 25 µm (**D**) Quantification of the average length of the AIS (**E**) Quantification of the distance of the AIS from soma. For both D and E: *, p<0.05; **, p<0.001; one-way ANOVA; n=4, mean ± SEM (**F**) Glutamatergic neurons were generated from human iPS cells. After 4 weeks maturation, the cells were treated with Aβ oligomers for 24 hours before fixation for immunostaining for Tuj1 (red) and AnkG (green). Scale bar = 10 µm. (**G**) Quantification of the AIS length (**H**) Quantification of the distance of the AIS from soma from the soma. (**I**) DIV7 neurons transfected with *APP*_*Swe*_ plasmid with or without the calpain inhibitor, PD150606 (PD), for two days and immunolabeled with the AIS marker βIV spectrin (yellow) and APP (6E10, red). GFP is shown with green. Scale bar = 10 µm. (**J**) Quantification of the length of the AIS from the experiment shown in panel I. (*, p<0.05; **, p<0.001; one-way ANOVA; n=4; Data represent means ± SEM.)

To determine whether the mouse neurons we were using for our Aβ assays accurately reflected the situation in the human brain, we established iPSC-derived neuronal cultures to test the effect of Aβ on the dimensions of the AIS of human neurons. Four to five weeks after induction (see Methods), the mature human iPSC neurons were treated with Aβ(o) for 24 h followed by immunostaining for the neuronal marker, TuJ1, and the AIS marker, AnkG (Fig. 3F). In untreated cultures the average length of the AIS in the human neurons (∼30 µm) was similar to the length of the AIS in cultured mouse neurons (∼35 µm). The distance from the cell soma to the most proximal portion of the AIS was slightly greater in human iPS neurons (∼15 µm) compared with mouse primary neurons (∼11 µm), but remarkably similar given the differences in the two preparations. The similarities carried over into the Aβ response. Neither the length of the AIS (Fig. 3G) nor its distance from the soma (Fig. 3H) was altered in the human neurons by the application of Aβ. These data suggest that the alterations of the AIS are caused by the overexpression of APP directly and not indirectly through increased levels of Aβ.

The AD brain has evidence of increased calpain protease activity (Saito et al., 1993; Sheehan et al., 1997; Leissring et al., 2000), as do its mouse models (Kurbatskaya et al., 2016; Ahmad et al., 2018). Both AnkG and βIV spectrin are calpain substrates and their proteolytic cleavage leads to the dispersion of the ion channels normally clustered in the AIS (Schafer et al., 2009; Greer et al., 2013). Glutamate stimulation would be expected to activate NMDA channels, increase intracellular calcium, and thus potentially activate calpain. APP has also been reported to alter calcium dynamics, although there are questions as to how and whether this happens (Woods and Padmanabhan, 2012). To test whether the reduction of AIS length we observed was due to the activation of calpain, we pretreated primary cultures of wild type mouse neurons with the calpain inhibitor, PD150606 (1 mM), and transfected the neurons with APP_Swe_. After calpain inhibition, we found that the reduction in AIS length was significantly less than in cells without inhibitor (Fig. 3I, J). Immunocytochemistry with 6E10 showed that the same amount of APP was expressed and its distribution within the cells was comparable in the presence and absence of drug. It should be noted, however, that the length of the AIS was only partially restored by this treatment (Fig. 3I). The suggestion is that while calpain plays a role in the APP effects, there are other mechanisms involved.

#### The AIS in Alzheimer’s disease – in vivo data

The data described thus far have come from in vitro assays. To ask whether similar results would be found in vivo, we first compared the AIS in cryostat sections from the brains of wild type and R1.40 mice. Recall that R1.40 mice carry in their genome a human *APP*_*Swe*_ transgene in addition to the endogenous (wild type) mouse *App* gene. In wild type brains, the AIS in vivo was considerably shorter (10-15 µm) than found in the in vitro cortical cultures of either human or mouse neurons. Even with this difference, at 18 months of age the average AIS of R1.40 neurons in vivo was significantly shorter than those of wild type neurons (Fig. 4A). The reduction was seen in all regions of cortex but failed to reach statistical significance in posterior (visual) cortex (Fig. 4B). These measurements were all made in sections taken in the sagittal plane. To rule out the possibility that a change in orientation rather than in length was the true cause of this difference, we also took AIS measurements of cortical neurons in the coronal plane and obtained similar results (Fig. 4C). These changes in length were found even though the density of the AIS did not differ between the two genotypes (Fig. 4D, E). We also quantified the AIS lengths of pyramidal cells of the CA fields of the hippocampus (Fig. 4F, H) and found them to be significantly shorter in both the sagittal (Fig. 4F) and coronal planes (not shown). Unlike the neocortex, however, the density of AIS was significantly reduced in the CA fields (Fig. 4G). We did not analyze the neurons of the dentate gyrus due to the intertwined nature of the AIS in this region.

**Figure 4.**
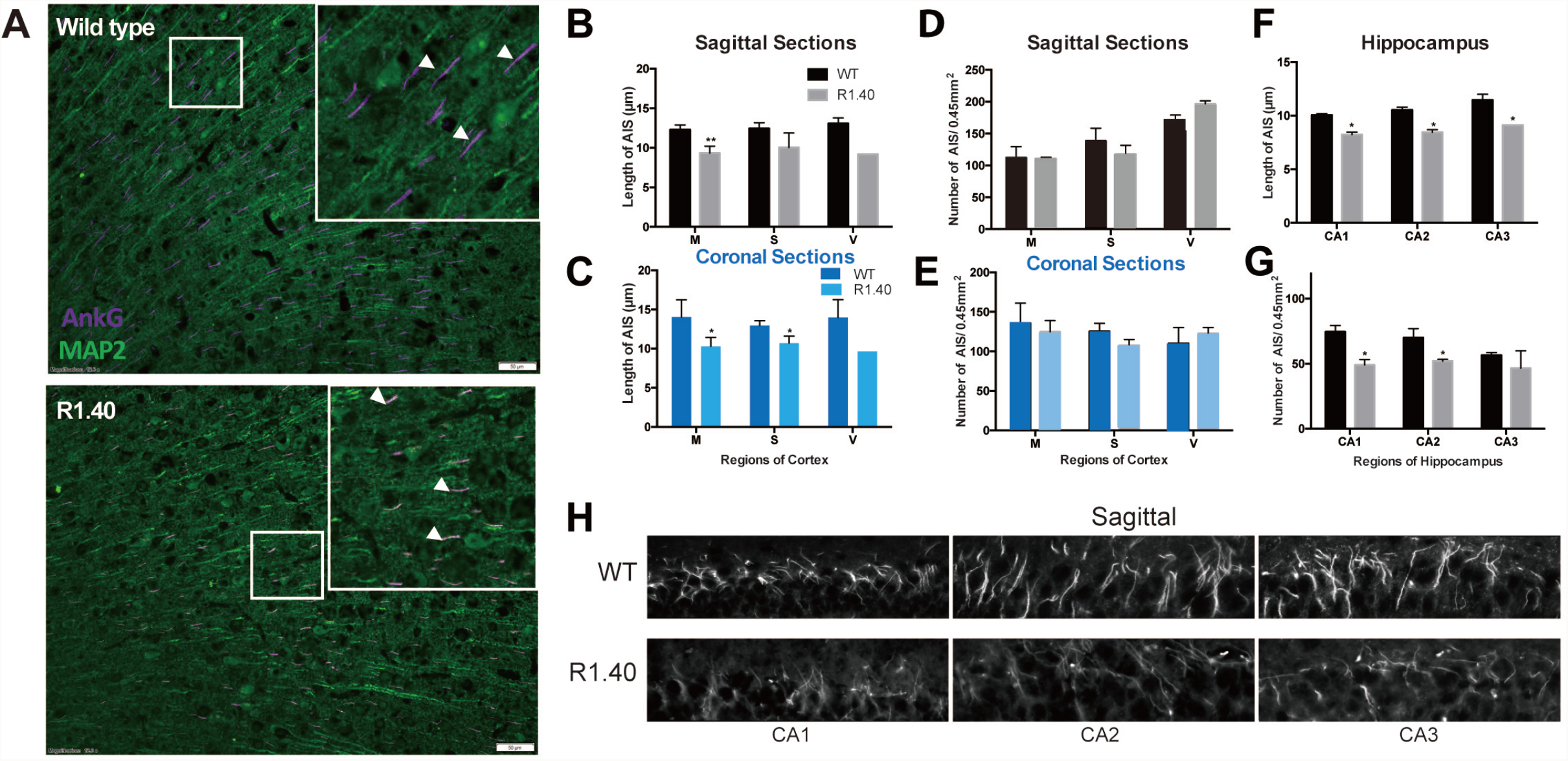
Alterations in the AIS in R1.40 cortex and hippocampus. (**A**) Cryostat sections from 18-month-old wild type and R1.40 frontal cortex immunostained with MAP2 (green) and AnkG (magenta) antibodies. Scale bar = 50 µm. (**B**) Sagittal sections from motor (M), somatosensory (S) and visual (V) cortex in 18-month wild type and R1.40 mice (**C**) Coronal sections from in 18-month wild type and R1.40 mice (**D**) AIS density (number of AIS/ 0.45 mm^2^ field) in motor (M), somatosensory (S) and visual (V) cortex measured in sagittal sections (**E**) AIS density in motor (M), somatosensory (S) and visual (V) cortex measured in coronal sections. (**F**) Length of the AIS of pyramidal neurons of the hippocampus of wild type and R1.40 mice observed in sagittal section (**G**) Density of the AIS of CA1, CA2, and CA3 pyramidal neurons of the hippocampus of wild type and R1.40 mice observed in sagittal sections. (**H**) Immunolabeling of the AIS of hippocampal CA1, CA2, and CA3 pyramidal cells in wild type and R1.40 mice. Scale bar = 10 µm. (*, p<0.05; **, p<0.001; unpaired t-test; n=4. Data represent means ± SEM.)

The in vitro data demonstrate that exposure to Aβ peptides alone does not affect the properties of the AIS. Qualitative observations of β4 spectrin/6E10 double-labeled material supported this view as they revealed no obvious bending of the AIS in or near cortical amyloid plaques (Fig. 5A). To provide quantitative detail to observations such as these we measured the density and length of the AIS in three concentric rings centered on 6E10-positive Aβ plaques in 18-month R1.40 mouse frontal cortex (Fig. 5B), a method established by Marin and Rasband (Marin et al., 2016). We found no statistically significant association between AIS length (Fig. 5C) or density (Fig. 5D) and the distance of the AIS from the center of a plaque. Thus, consistent with our in vitro findings, the reduction of AIS length was not sensitive to the presence of amyloid.

**Figure 5.**
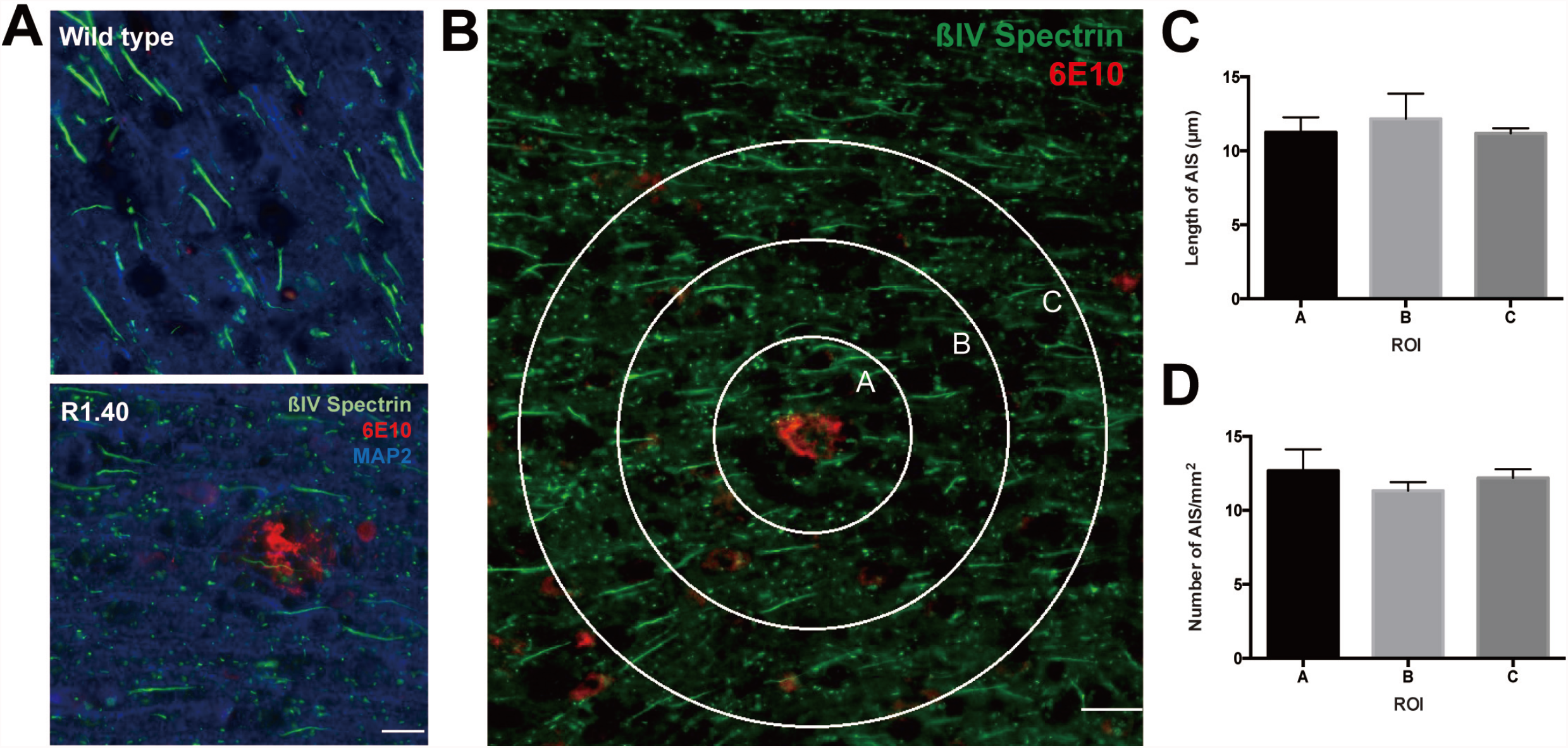
Properties of the AIS are unchanged in the vicinity of a plaque. (**A**) Cortex of 18-month wild type and R1.40 mouse labeled for βIV spectrin (green), APP (6E10 antibody, red) and MAP2 (blue). Scale bar = 10 µm. (**B**) Three concentric rings (50, 100, and 150 µm in diameter) were centered on an Aβ (6E10+) plaque and used to define three regions of interest (A, B, and C). AIS were immunostained using antibodies against βIV spectrin (green). Scale bar = 20 µm. (**C**) Quantification of the length of the AIS in the three regions. (**D**) Quantification of the distance of the AIS from the cell soma. The differences shown in panels C and D were not significant by one-way ANOVA; n=3. Data represent means ± SEM.

To test whether the situation in human Alzheimer’s disease was similar to the R1.40 mouse AD model, we turned to human post-mortem brain samples. We chose prefrontal cortex (Brodmann Area 9 – BA9) as our test region. We immunostained 10 µm cryostat sections for MAP2, APP (N-terminus), and AnkG with a DAPI nuclear counterstain. In material from healthy controls (Fig. 6A), the AIS were well stained, and many (though not all) could be clearly associated with their neuron of origin (Fig. 6Ai). While in control specimens the levels of intracellular APP immunoreactivity were low, a few neurons displayed elevated staining (yellow arrow, Fig. 6A). In cells with low levels of APP an associated AIS could often be identified (Fig. 6Aii). For cells with elevated APP, however, we could rarely identify its AIS. To be sure, the thickness of the cryostat sections was less than the average diameter of a neuron. Thus, for any given neuron, the AIS could well have been located in an adjacent section. The consistency of the finding, however, was striking and it was amplified by the situation in the AD material.

**Figure 6.**
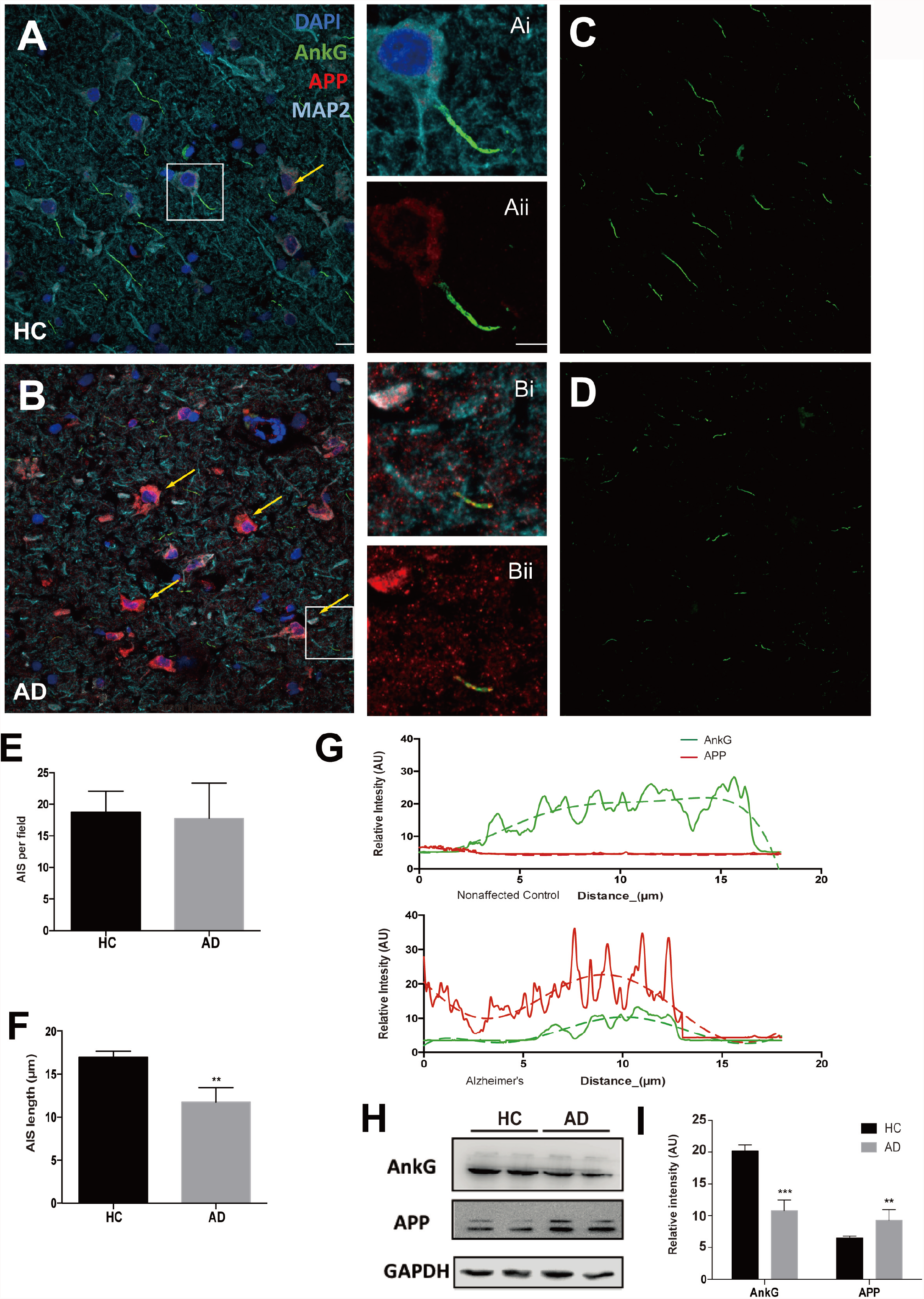
The impact of sporadic Alzheimer’s disease on the properties of the AIS. (**A**) Immunolabeling of the AIS in neurons from Brodmann Area 9 (BA9) of a healthy control (HC) using antibodies against AnkG (green), N’APP (red), MAP2 (cyan), and DAPI (blue). Yellow arrows indicate neurons with increased intracellular APP. (**Ai**) Enlargement of the area shown in the white box; (**Aii**) Same area as (Ai) showing only the APP and AnkG labels. (**B**) Immunolabeling of the AIS in neurons from BA9 of a case of sporadic Alzheimer’s disease (AD). Yellow arrows indicate neurons with increased intracellular APP. (**Bi**) Enlargement of the area shown in the white box; (**Bii**) Same area as (Bi) showing only the APP and AnkG labels. (**C**) Same field as (A) showing only the AnkG channel. (**D**) Same field as (B) showing only the AnkG only channel. Scale bars = 10 µm. (**E**) The density of AIS in the BA9 region of healthy controls and sporadic AD. (**F**) The length of the AIS in the BA9 region of healthy controls and sporadic AD. (**G**) Intensity of AnkG immunolabeling along the AIS show in panels Aii and Bii (AnkG, green lines and APP, red lines). Green and red dashed lines represent the polynomial regression curve of the intensity of AnkG and APP, respectively. (**H**) Western blots of BA9 lysates of age-matched HC and AD brains probed with AnkG and APP antibodies. (**I**) Quantification of the band intensity of the western blot data shown in panel (H). **, p<0.001; unpaired t-test; n=6. Data represent means ± SEM.

In all six AD cases examined the presence of amyloid plaques was readily observed. Though less often commented on in the literature, we found an equally impressive increase in the level of background staining. This background can be clearly seen in the insets to Figure 6B (Fig. 6Bi, 6Bii). As each of the AD cases was sporadic in nature the elevated levels of APP immunostaining were most probably a function of age and environmental factors. In addition to the higher APP background, virtually every neuron in cortex presented with strong intracellular APP immunoreactivity (yellow arrows, Fig. 6B). Plus, as would be predicted from the few high APP expressing neurons in the healthy controls, we could almost never define a cell of origin for any of the AIS present in an AD brain section.

The AnkG channel from the merged images (Fig. 6A, B) was used to quantify the dimensions of the AIS in healthy controls (Fig. 6C) and in AD cases (Fig. 6D). It was notable that, despite the fact that the human brain is several thousand-fold larger than the mouse brain, the length of the AIS of human prefrontal neurons (10-18 µm) was only slightly longer than that found in the mouse (10-15 µm). By contrast, the average density of AIS in the human material was about a fifth of that found in mouse (Fig. 6E). Further, just as we found in the R1.40 transgenic mouse, the average length of the AIS was decreased by about one-third in the Alzheimer’s disease brain compared to healthy controls (Fig. 6F). Double-staining with AnkG and N-term APP plainly showed that in all AD cases the APP protein had infiltrated the AIS region (Fig. 6Bii). An important distinction with the mouse, however, was that there was little evidence of a reciprocal gradient of APP and AnkG. In the R1.40 mouse, the excess APP appeared to shift the peak of AnkG nearly 10 µm distally from that seen in wild type. In the human, even though the intensity and total length of AnkG staining region was reduced the shift was less apparent (Fig. 6G, compare with green symbols in Fig. 2I).

To obtain a more global picture of these changes we performed western blots of APP and Ankyrin G using lysates prepared from unused cryostat sections of the same blocks of human BA9 tissue (Fig.6H). Not unexpectedly, we found that the levels of APP protein were considerably higher on average in the AD brain than in healthy controls (2-3-fold, Fig 6I). The increase was statistically significant (n = 6 healthy controls and 6 AD cases) and quantitatively similar to the increase seen in APP levels in the R1.40 mouse brain. By contrast, the overall levels of Ankyrin G protein were significantly decreased in AD brain compared to healthy controls (Fig. 6I).

## Discussion

The findings here reveal several new and unexpected roles of the full-length APP protein in brain function. First, we found that a sustained increase in neuronal activity leads to a rapid increase in *App* gene expression. Second, we demonstrated that elevated levels of APP protein – whether caused by neuronal firing, APP overexpression, or an APP transgene – lead to two important changes in the axon initial segment (AIS): it shortens in length and shifts in position away from the cell body. Third, we showed that the effect on AIS length is observed in mouse neurons both in vivo and in vitro. Finally, we confirmed that in persons with sporadic Alzheimer’s disease (AD) cortical neurons have elevated levels of APP protein, and that portions of this excess APP invade the AIS domain. This increased APP is associated with a significant shortening of the AIS compared to healthy controls. All four of these findings have significant implications for the understanding of AD.

The AIS is an important yet highly dynamic region of a neuron. Alterations of the length and position of the AIS are activity-dependent and, in turn, affect neuronal activity (Grubb and Burrone, 2010a; Huang and Rasband, 2018; Leterrier, 2018). As the site of action potential initiation, even subtle changes in the AIS can have measurable effects on neuronal excitability (Grubb and Burrone, 2010c, b; Kuba et al., 2010; Kuba, 2012; Atapour and Rosa, 2017; Kneynsberg and Kanaan, 2017). A longer AIS, or one more proximal to the cell body, will result in a more excitable neuron due to a lower action potential firing threshold. The reverse situation is also true: when the length of the AIS decreases or it shifts to a more distal position along the axon, neuronal activity is decreased (Grubb and Burrone, 2010b; Rasband, 2010). While this correlation between AIS structure and electrophysiological function has been clear, the mechanisms that link changes in AIS structure with activity have remained largely unknown. Correlations with altered calcium levels and the resulting activation of calpain, a protease capable of cleaving both Ankyrin G and βIV spectrin, offer one possible explanation. Changes in the natural calpain inhibitor, calcineurin (Evans et al., 2013a) support this hypothesis. Yet the studies described here using calpain inhibition suggest that calpain can provide only a partial explanation of the observed phenomena.

The actions of APP we have documented offer a compelling additional mechanism to explain the correlation between neuronal activity and AIS behavior. With immunocytochemistry in both mouse and human we showed that, in a mature AIS, APP is in close apposition with both ankyrin G and βIV spectrin. The results of the co-immunoprecipitation experiments confirm this by showing that APP physically interacts with both major scaffolding proteins. The association is found with human as well as with mouse APP proteins. Wild type mice produce exclusively mouse APP while the R1.40 animals produce mouse APP plus 2-3 times that amount of human APP (hAPP_Swe_). Manipulating the levels of APP either genetically or through chronic glutamate stimulation shows that this physical interaction is coupled with structural consequences for the AIS. With increased neuronal activity, *App* gene transcription is rapidly upregulated, and protein levels follow rather closely. The transcriptional response is mostly likely the key to this regulation as the translated APP protein has a relatively short half-life (Xiao et al., 2015). Coupled with the electrophysiological consequences of changes in AIS structure, the dynamic nature of the transcriptional response suggests that under normal conditions APP acts as a “governor” on neuronal excitability. When neuronal activity goes up, APP levels rise causing the distance between the AIS and the soma to increase and the length of the AIS to decrease. The overall effect is to reduce the neuron’s firing threshold and thus serve as a protective mechanism.

The data from neurons in human AD and its mouse models suggests that beyond its contribution to normal neuronal activity, the AIS may also play a role in neurodegenerative disease. Elevated APP levels such as those found in Alzheimer’s drive the same changes in AIS position and length as does increased neuronal activity. The relevance of this finding to AD is enhanced by data showing that hAPP_Swe_ has effects on the AIS that are more dramatic those of wild type hAPP. The effects are cell autonomous, caused by endogenous full-length APP protein, not by exogenous Aβ. Addition of Aβ to the culture medium (oligomers or fibrils) has no effect on the AIS of either human or mouse neurons. Plus, the length of the AIS in the neurons of the mouse brain is unaffected by its proximity to an amyloid deposit. The immunoprecipitation data further support the connection between the dynamics of the AIS and the symptoms of AD. Antibody to APP pulls down ankyrin G and βIV spectrin from both wild type and R1.40 brain lysates, but the interaction appears stronger in R1.40 than in wild type brain lysates. Even the retarded development of the AIS in R1.40 cultures may have implications for the full clinical picture of Alzheimer’s. AD has long been recognized as having a developmental component, with intellectual performance in youth inversely related to the risk of dementia in old age (Snowdon et al., 1996; Whalley et al., 2000). In this light, the slow development and shorter length of the AIS in cultured R1.40 neurons suggests that elevated APP expression may be one explanation for this connection.

In sporadic Alzheimer’s disease we speculate that the aging process leads to a relentless increase in the damage experienced by neurons. As APP is a damage response protein this results in a chronic elevation in the levels of APP. This lifetime accumulation of APP, worsened by hyperexcitability, irritation due to inflammation, vascular problems or a traumatic injury eventually tips the system into a disease state. The levels of APP never return to normal leading to a sustained shift in the position of the AIS away from the cell body and a reduction in its length. Both changes would have the effect of lowering neuronal excitability. We found that the reduction of AIS length compared to control animals was uniformly distributed throughout the R1.40 frontal cortex. This is a direct prediction of our model as it is APP-dependent, not Aβ-dependent. The effects on the AIS can be assumed to occur throughout the nervous system, not only in proximity to amyloid plaques. Our model further predicts that familial AD mutations in APP exacerbate the normal age-associated vulnerabilities, leading to an earlier aging of onset, as is seen.

One important observation that does not fit our proposed model is that hyperexcitability and epileptic activity is a feature of both Alzheimer’s disease and its mouse models (Di Lazzaro et al., 2004b; Di Lazzaro et al., 2004a; Palop et al., 2007; Minkeviciene et al., 2009; Pennisi et al., 2011), yet one of the phenotypes of APP overexpressing neurons is a reduced probability of firing. Though only speculation, we would propose that the APP over-expression and its effects on the AIS are not experienced equally by all brain regions and that those regions, or neurons, that are spared attempt to compensate for reduced network activity increasing their firing frequency. This is consistent with the heterogeneity we observed in our human material in both the APP intensity in different neurons and the infiltration of APP in different individual AIS.

One of the strongest pieces of evidence supporting the amyloid cascade hypothesis has been the identification of APP as an autosomal dominant Alzheimer’s disease gene. Formally, however, the genetics point only to APP, not to Aβ. While the discovery of the *PSEN1* and *PSEN2* Alzheimer’s disease genes could also be reasonably argued to fortify the Aβ argument, this is only indirect support as the γ-secretase has dozens of substrates apart from APP. These doubts are magnified by the realization that, despite many years of investigation, the molecular mechanisms by which the Aβ peptide kills a neuron remain only speculative. Further, the areas of the AD brain where neurodegeneration is most evident are not the ones with the highest density of Aβ plaques. We propose that the actions of the full-length APP protein on the AIS and the resultant disruption of network activity provide an alternative explanation for the APP genetic data. In this conceptualization, the effect of APP on the pathogenesis of Alzheimer’s disease is through its action on the AIS, not through the generation of Aβ.

